# Delineating the shape of COPII coated membrane bud

**DOI:** 10.1101/2024.02.13.580145

**Authors:** Sanjoy Paul, Anjon Audhya, Qiang Cui

## Abstract

Curvature-generating proteins that direct membrane trafficking assemble on the surface of lipid bilayers to bud transport intermediates, which move protein and lipid cargoes from one cellular compartment to another. Our recent study on the COPII protein Sar1 showed that the inserted volume of the protein into the membrane determines the degree of membrane bending. However, it is unclear what controls the overall shape of the membrane bud once curvature induction has begun. In vitro experiments showed that excessive concentrations of Sar1 promoted the formation of membrane tubules from synthetic vesicles, while COPII-coated transport intermediates in cells are generally more spherical or lobed in shape. To understand the origin of these morphological dissimilarities, we employ atomistic, coarse-grained (CG), and continuum mesoscopic simulations of membranes in the presence of multiple curvature-generating proteins. We first demonstrate the membrane bending ability of amphipathic peptides derived from the amino terminus of Sar1, as a function of inter-peptide angle and concentration using an atomistic bicelle simulation protocol. Then, we employ CG (MARTINI) simulations to reveal that Sec23 and Sec24 control the relative spacing between Sar1 protomers and form the inner-coat unit through an attachment with Sar1. Finally, using Dynamical Triangulated Surface (DTS) simulations based on the Helfrich Hamiltonian we demonstrate that the uniform distribution of spacer molecules among curvature-generating proteins is crucial to the spherical budding of the membrane. Overall, we show that Sec23 and Sec24 act as a spacer to preserve a dispersed arrangement of Sar1 protomers and to help determine the overall shape of the membrane bud.

## Introduction

COat Protein complex II (COPII) is a multiprotein molecular machinery that orchestrates the export of newly synthesized proteins from the Endoplasmic Reticulum (ER) via membrane-enclosed transport carriers.(1) The complex consists of various isoforms of Sar1, Sec23, Sec24, Sec13, and Sec31, which are thought to assemble into a multilayered coat structure on the cytoplasmic face of discrete ER subdomains known as transitional ER. Sar1 initiates the membrane budding process when it becomes activated by transitioning from a GDP-bound state to a GTP-bound state by the guanine nucleotide exchange factor Sec12.(2; 3) Subsequently, Sar1 together with Sec23-Sec24 heterodimers forms the inner coat layer(4; 5; 6) on the membrane bud, whereas Sec13-Sec31 produces an outer cage-like layer(7; 8) to complete COPII coat formation. These membrane-bound, cargo laden carriers generally adopt a spherical or multi-lobed shape that are roughly ∼ 50-200 *nm* in diameter.(9) With the help of Sec16 and members of the TANGO1 family, multiple COPII coated carriers can adopt a ‘beads-on-a-string’ conformation to accommodate bulky procollagens.(10; 11) However, in-vitro experiments have demonstrated that Sar1 in the presence of GTP forms an organized lattice structure on Giant Unilamellar Vesicles (GUVs), resulting in the formation of membrane tubules.(12; 13) This shape is noticeably different from the structure of COPII-coated transport carriers found inside living organisms. Further, this indicates that Sar1 alone cannot produce the spherical shape of a membrane bud. Therefore, the molecular origin of the shape of COPII coated carriers remains poorly understood.

Regulation of the spatiotemporal accumulation of COPII proteins is crucial to drive cargo export from the ER. Our recent study highlighted the formation of the inner-coat layer as the rate-limiting step for the cargo transport process.(14) We also demonstrated the molecular mechanism of membrane binding and bending activity of inner coat protein Sar1 in a nucleotide state- and concentration-dependent manner.(15) Despite these advances, what regulates the shape of membrane buds induced by COPII needs to be understood in more detail. Conventionally, it is considered that Sec23(16) functions as a GTPase activating protein (GAP), facilitating the hydrolysis of GTP on Sar1, whereas Sec24(17) is involved in cargo binding. However, in the absence of Sar1, cargo export persists through the continued action of Sec23-Sec24 in a phase separated state.(18) These findings suggest that Sec23-Sec24 complexes may play hitherto unrecognized roles in the process of membrane budding.

Sar1 inserts its amphipathic amino terminal helix into the membrane to induce local positive curvature.(15; 19) While a single Sar1 protein can locally deform the membrane, how multiple Sar1 molecules sculpt the membrane into distinct shapes remains unknown. The consequence of multiple curvature-inducing inclusions on membrane has been shown to produce a diverse array of morphologies such as tubes, corkscrew, disc, caveolae etc.(20; 21) In the context of COPII mediated membrane budding, relevant membrane shapes are tubules under *in vitro* conditions and more spherical under *in vivo* conditions. To comprehend the topology of the membrane bud formed by COPII, it is crucial to establish how the membrane responds to the multiple types of proteins involved.

In this study, we employ atomistic, coarse-grained, and continuum mechanics based simulations to elucidate the molecular mechanism of how the inner coat layer shapes the membrane bud. Since the amphipathic amino terminal helix is the curvature-inducing region of Sar1, we arrange multiple such peptides on the membrane and study their collective membrane bending activities. First, we compare the relative bending activities of a GTP bound Sar1 dimer vs. the amino terminal peptide dimer in the absence of the rest of the protein using the atomistic bicelle simulation protocol. Then, we provide a quantitative estimate of the magnitude of curvature induction as a function of relative orientation of the amino terminal peptides and their concentration. We also investigate the membrane binding ability of Sec23 and Sec24 using MARTINI based coarse-grained model. Finally, we utilize the Dynamic Triangulated Surface (DTS) simulation framework to explore the relationship between the surface coverage of Sar1 and the shape of the membrane bud. Taken together, our model suggests a new role for Sec23,Sec24, and cargo proteins in COPII mediated membrane budding process in which they participate as spacers to control the relative distribution of Sar1 proteins on the lipid bilayer surface, which is critical for the spherical development of the membrane bud.

## Results

### Comparative assessment of the membrane bending activity of the amino terminal amphipathic helix of Sar1 in the presence and absence of the rest of the protein

We first investigate the membrane bending activity by amphipathic helices when present in isolation vs. accompanied by the rest of the protein. GTP bound Sar1 dimer (h-GTP dimer) produces significant positive curvature on the membrane, transforming the flat-shaped bicelle (Fig-1A) into a highly bent dome-shaped structure (Fig-1C). The height of phosphate mid-plane changes by ∼ 4 nm as a consequence of this transition. However, when the amphipathic (Fig-1D) helices are present alone (arrangement-1) without the rest of the protein, the magnitude of curvature induction is dramatically reduced (Fig-1E-F). Although the protein segment that causes curvature induction is the same for both cases, we observe a stark contrast in the extent of membrane bending.

**Figure 1.**
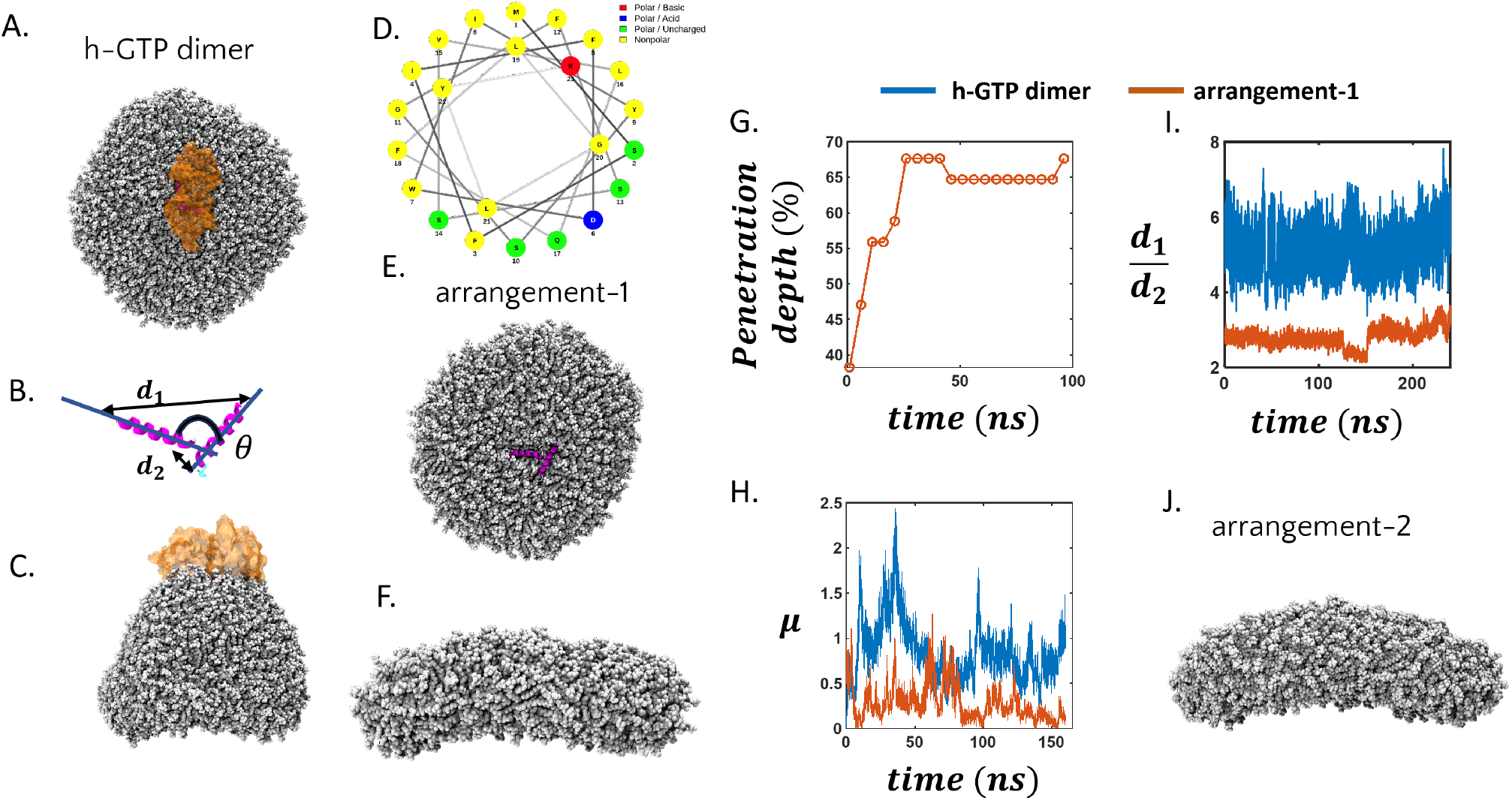
Isolated amphipathic regions of Sar1 diminish its capacity to bend the membrane. (A) Top view of h-GTP dimer protein on membrane bicelle. The amphipathic helix is coloured in purple. The protein is depicted in surface representation with orange colour and the membrane is shown in VDW represntation with silver colour. (B) Arrangement of the helix-pair (residue 1-23) derived from the amino terminal amphipathic region in h-GTP dimer. End distances of the peptide pair is represented by *d*1 and *d*2 and the inter-peptide angle is *θ* (C) Side view of the h-GTP dimer membrane system after ∼248 *ns* simulation revealing a highly bent shaped membrane as a consequence of high curvature induction. (D) Helical wheel diagram of the amino terminal amphipathic helix of h-GTP Sar1 where different colours represent different residue types (yellow: hydrophobic, green: polar uncharged, blue: polar acidic and red: polar basic) (E) Top view of the membrane bicelle system with peptide-arrangement-1 (isolated amphipathic peptide case) (F) Snapshot of the membrane bicelle with peptide-arrangement-1 after 250 *ns* of simulation. Time evolution of (G) membrane penetration depth and (H) *µ* for the h-GTP dimer and peptide-arrangement-1 in the presence of continuous membrane. (I) *d*1*/d*2 as a function of time in case of membrane bicelle simulation in the presence of h-GTP dimer and peptide-arrangment-1. (J) Snapshot of the membrane bicelle system in the presence of peptide-arrangemnt-2 where *θ* is restrained to 180°.

To explain this differential bending activity of the Sar1 amphipathic helices, we study the time evolution of the penetration depth of the peptides into a periodically continuous membrane. While the h-GTP dimer exhibits 40 % penetration depth (as defined by Paul et. al.(15)), the amphipathic helix increases the penetration depth up to 65 % within ∼100 *ns* when present in isolation (Fig-1G). This deeper membrane penetration in the absence of the rest of the protein reduces the magnitude of partitioning of hydrophobic/hydrophilic residues at the membrane-water interface. While the h-GTP dimer displays *µ*∼1 (Eqn. 1), the isolated peptides (arrangement-1) lead to *µ* significantly less than 1 (Fig-1H). This indicates that in the absence of the rest of the protein, the amphipathic helix inserts into the membrane so deeply that it brings some of its hydrophilic residues into the membrane. As a result, the inter-leaflet stress, which arises due to the hydrophobic/hydrophilic partitioning at the membrane water interface, becomes reduced and therefore does not bend the membrane significantly.

In addition to the excess penetration of the peptides, another contributing factor in this context can be the relative orientation of the peptides on the membrane surface (Fig-1B). The distribution of the inter-peptide angle (*θ*) in the h-GTP dimer is sharply peaked around ∼105° whereas the peptides in isolation following arrangement-1 exhibit a broad distribution of *θ* (Fig-S1A). We further characterize this orientational dissimilarity by computing the ratio of the end distances (*d*1 and *d*2) of the peptide pair. In the case of a serial arrangement of the peptides *d*1*/d*2 will be 2*d* where *d* is the length of a single helix (∼3.5 *nm* in the case of Sar1), and in the case of a parallel arrangement, *d*1*/d*2 should be 1. The high value of *d*1*/d*2 observed for the h-GTP dimer indicates a serial-like arrangement of the helix-pair while the significantly lower value of *d*1*/d*2 for the peptides in isolation indicates rather different arrangements (Fig-1I). To estimate the effect of the inter-helix orientation on the membrane curvature induction, we perform bicelle simulations in the presence of two peptides while restraining the inter-peptide angle to be 180° (peptide-arrangement-2). The angular restraint is not observed to enhance the degree of membrane bending (Fig-1J). Therefore, the relative orientation of the amphipathic peptides does not significantly impact the membrane bending activity.

This suggests that curvature induction by a large assembly of the amphipathic peptides can be modeled without considering specific relative orientation (*vide infra*).

### The magnitude of membrane curvature induction is proportional to the concentration of the embedded peptides

In this section, we study the impact of the peptide concentration on the magnitude of membrane curvature induction. From the previous section, it is clear that two copies of the amphipathic peptide derived from the h-GTP dimer are unable to generate significant membrane curvature irrespective of their relative orientation. With one more amphipathic helix (peptide-arrangement-3) as shown in Fig-2A, the extent of membrane bending increases from ∼ 0.01 *nm*^−1^ (arrangement-1 and 2) to 0.013 *nm*^−1^

**Figure 2.**
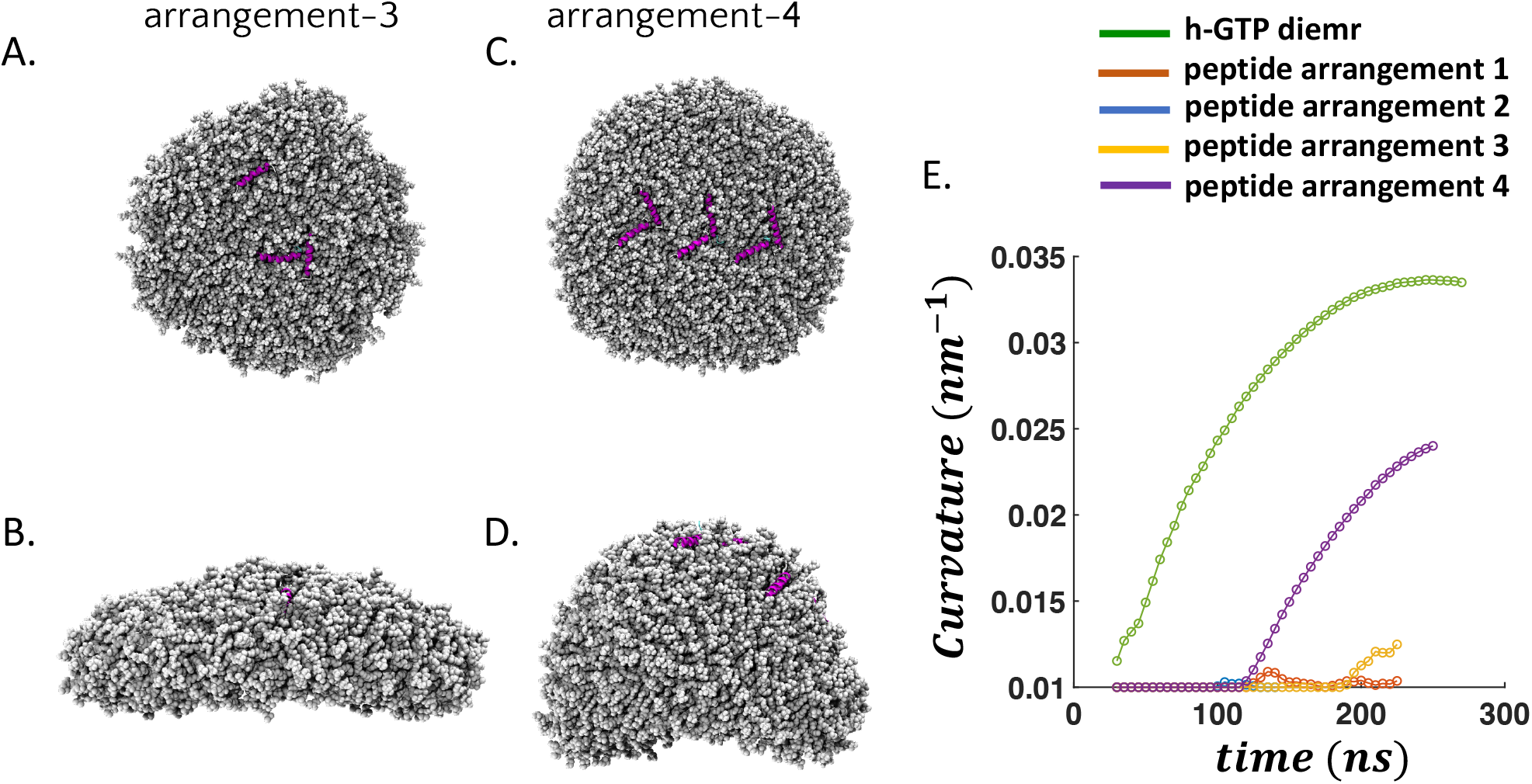
Curvature induction on the membrane as a function of peptide concentration. (A) Initial (top) and (B) final (side) snapshot of a tri-peptide assembly (peptide-arrangement-3) on a membrane bicelle after ∼ 250 *ns* simulation. (C-D) Initial and final snapshot in case peptide arrangement 4. (E) Time evolution of membrane curvature in all the cases of peptide arrangements and h-GTP dimer protein. h-GTP dimer protein exhibit highest curvature followed by peptide-arrangement-4. Peptide arrangement 1-2 show lowest curvature induction.

(Fig-2B). With 6 peptides (arrangement-4) distributed around the center of the bicelle (Fig-2C), we observe strong bending of the membrane, leading to a vesicular cap like structure similar to induced by the h-GTP dimer (Fig-2D) with a curvature of ∼ 0.25 *nm*^−1^ Fig-2E). Thus, the bicelle simulations demonstrate that peptide assemblies induce membrane curvature in a concentration-dependent manner. Analysis of inter peptide angles reveals a broad distribution (50 − 180°), further confirming the lack of any strong correlation between membrane bending activity and inter-peptide orientation (Fig-S1 B-C). As a control simulation, we study a bicelle system covered with a total of 18 peptides (Fig-S2). Surprisingly, in this case, we do not observe any significant membrane bending. By the end of ∼250 *ns* of simulation, many peptides are located at the highly curved edges of the bicelle, and they propagate the stress induced by hydrophobic insertion throughout the membrane. As a result, the net inter-leaflet stress is minimal and no curvature induction is observed.

### Sar1 serves as a tether connecting Sec23-Sec24 to create the inner-coat layer

We next study the binding of different protein components from the inner-coat layer to the membrane, moving towards a more realistic description of COPII. To address this question in a computationally effective manner we adopt the coarse-grained MARTINI models, which were successfully employed to model protein(22) and liquid droplet(23) mediated remodeling of membranes. Here, we first investigate the binding of Sec23 and Sec24 separately to the membrane. Both proteins detach from the membrane rather quickly (tens of nanoseconds) during the simulation despite initial placement on the membrane surface (Fig-3A-D). These observations are consistent with the model that Sar1 acts as an anchor to bring Sec23 and Sec24 to the membrane surface and generate the inner-coat layer. Indeed, in a simulation with a Sar1-Sec23-Sec24 trimer on the membrane (Fig-3E-F), Sec23 and Sec24 securely attach to Sar1, thereby maintaining their binding to the membrane and producing a cohesive unit. Subsequently, we attempt to simulate a more realistic inner-coat layer by increasing the number of repeating units of Sar1-Sec23-Sec24 from 1 to 2, 4 and 8. When the number of Sar1-Sec23-Sec24 trimer units is 2, we do not observe spontaneous protein-protein association among the trimer units due to dilution of the protein concentration (Fig-S3). However, as we increase the number of trimer units we observe more trimer-trimer associations. In the case of 8 trimer repeating units, we observe that a few Sar1 proteins stay out of the membrane plane as a consequence of crowding (Fig-3G). Overall, we observe a stable binding of the inner-coat components to the membrane where proteins are dynamically coupled with each other. We further evaluate the spacing between the Sar1 proteins in the presence and absence of Sec23 and Sec24 by measuring the distances (*d*_*amino*_) between their amino-terminal amphipathic segments (Fig-3H). In the absence of Sec23 and Sec24, the Sar1 tetramer exhibits strong inter-protein binding (Fig-S3) resulting in *d*_*amino*_ values in the range of 0−10 *nm*. In the presence of Sec23 and Sec24, *d*_*amino*_ increases significantly (∼8-30 *nm*). It is important to note that we consider only the structured part of Sec24 in our simulation. The amino terminal (resid 1-132) of Sec24 is intrinsically disordered and not included in our model; it is expected to further separate Sar1 protomers due to entropic repulsion. Thus, Sec23 and Sec24 serve as spacers in the inner-coat layer to separate Sar1 proteins from one another and prevent them from co-assembling with one another. In addition to Sec23 and Sec24, the bulky cargo proteins that are being packaged may also serve as spacers.(24) In the next section, we explore the consequence of including spacers between the curvature-generating proteins on the shape of the generated membrane bud.

**Figure 3.**
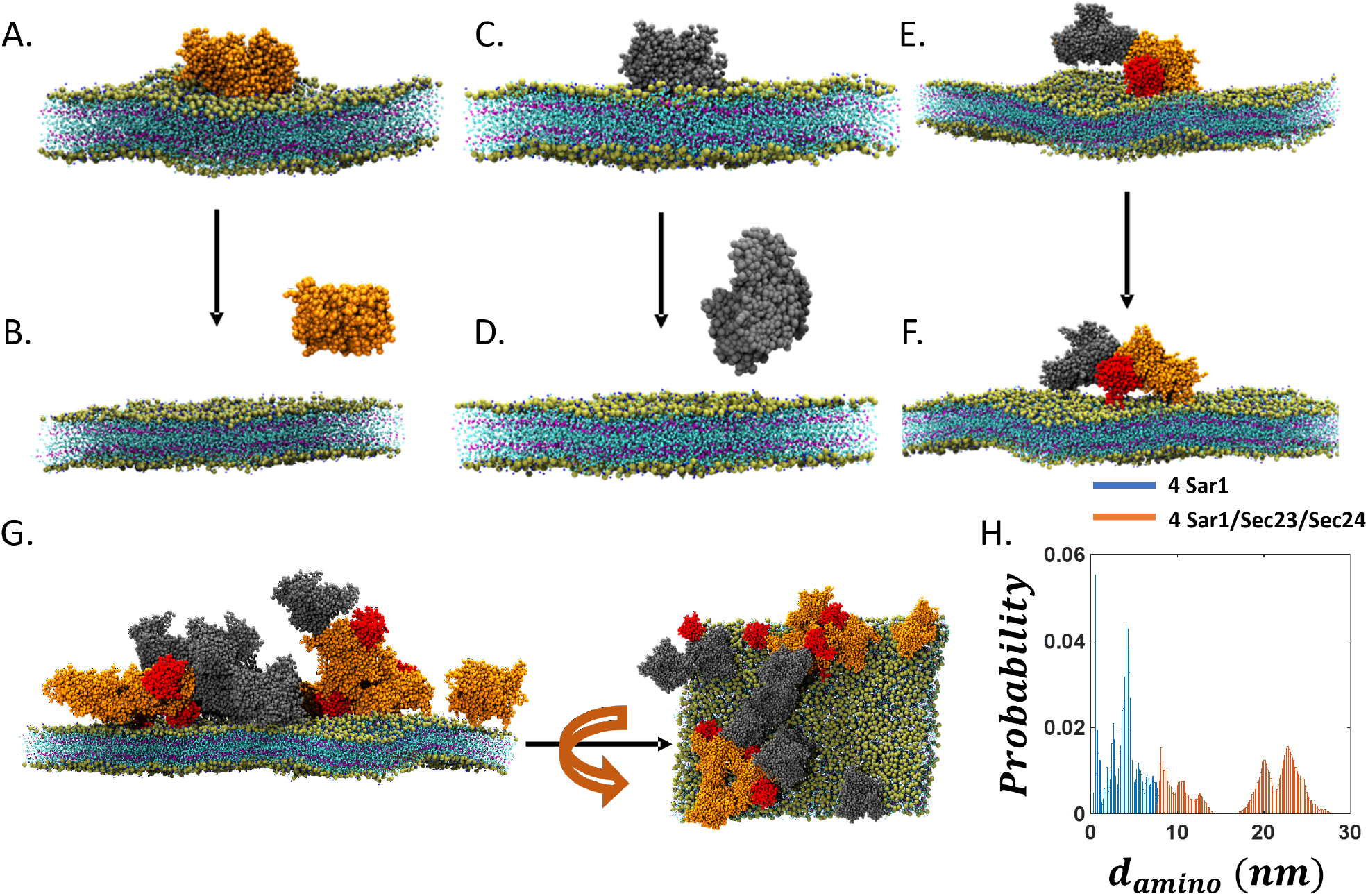
Simulating the inner coat layer using MARTINI based description. Initial (*t* = 0) and final (*t*∼2 *µs*) snapshots of (A-B) Sec23 (C-D) Sec24 and (E-F) Sar1-Sec23-Sec24 trimer in the presence of membrane. Protein coloured in red indicates Sar1 wheres Sec23 and Sec24 are depicted as orange and grey colour respectively. Sec23 and Sec24 individually fail to maintain a stable attachment with the membrane but with the help of Sar1 it remain bound to the membrane surface. (G) Top and side view of the 8 repeating units of Sar1-Sec23-Sec24 trimer. (H) Probability distribution of *d*_*amino*_ estimated from the simulations of Sar1 tetramer in the absence of Sec23 and Sec24 (cyan) and 4 trimer repeating units of Sar1-Sec23-Sec24 (brown).

### The effect of spacers on the shape of the membrane bud

Here, we describe how the spatial arrangement of curvature-generating proteins impacts the shape of the induced membrane bud employing the DTS simulation protocol. DTS simulations have been previously used to study the transformation of a membrane vesicle into tubes, discs, and other shapes when curvature induction takes place anisotropically.(20; 25; 26) With a flat membrane patch under constant tension, isotropic curvature-inducing inclusions have been shown to produce pearled tubule-like budding when the surface coverage of the proteins exceeds a certain threshold value.(27; 28) Using the atomistic bicelle simulations, we observe that isotropic curvature induction is the key characteristic in the case of Sar1-mediated membrane remodeling where the relative orientation of its amphipathic helices does not affect the magnitude of curvature induction. Results from the previous section also suggest that Sec23 and Sec24 maintain the spatial separation of Sar1 proteins. Based on these findings, we develop a mesoscopic model of the inner coat layer on a triangulated membrane mesh where protein-containing vertices (blue region) have positive intrinsic curvatures (*c*_0_ = 1.0 *d*^−1^) isotropically coupled to the Helfrich term. These curvature-inducing vertices represent Sar1 protomers while other vertices represent the membrane (*c*_0_ = 0 *d*^−1^). In this case, we observe tubular budding of the membrane with one or multiple tubules (Fig-4A). This condition resembles that in *in vitro* GUVs coated with Sar1, where tubular budding is commonly observed(12). A fraction of the protein containing vertices turns into a tubular shape while the rest remains flat surrounding the tubular region.

Next, to model the presence of Sec23 and Sec24, we include spacers (red) uniformly distributed in the protein-containing region. With 5% spacer, we observe a pearled tubule-like shape of the membrane bud (Fig-4B). The spacers occupy both tubular and flat protein-containing vertices. Due to the incorporation of spacers, only one tubule is generated in 4 trajectories and a small spherical budding is observed in traj-2. Upon increasing the fraction of spacers from 5 % to 15 %, the shape of the membrane bud becomes more spherical in nature (Fig-4C). A spherical shape is observed in almost all cases of membrane buds with a constricted neck. Only in the case of traj-2, a doubly pearled tubule is generated. Further increasing the spacer content to 25 % leads to a perfectly spherical membrane bud (Fig-4D). Thus, increasing the concentration of spacers alters tubular budding into a more spherical budding. When spacer content exceeds a certain threshold the membrane budding no longer takes place. While 50 % spacer leads to a reduced size of the buds, 75% spacer does not lead to any budding during the MC simulations (Fig-S4).

**Figure 4.**
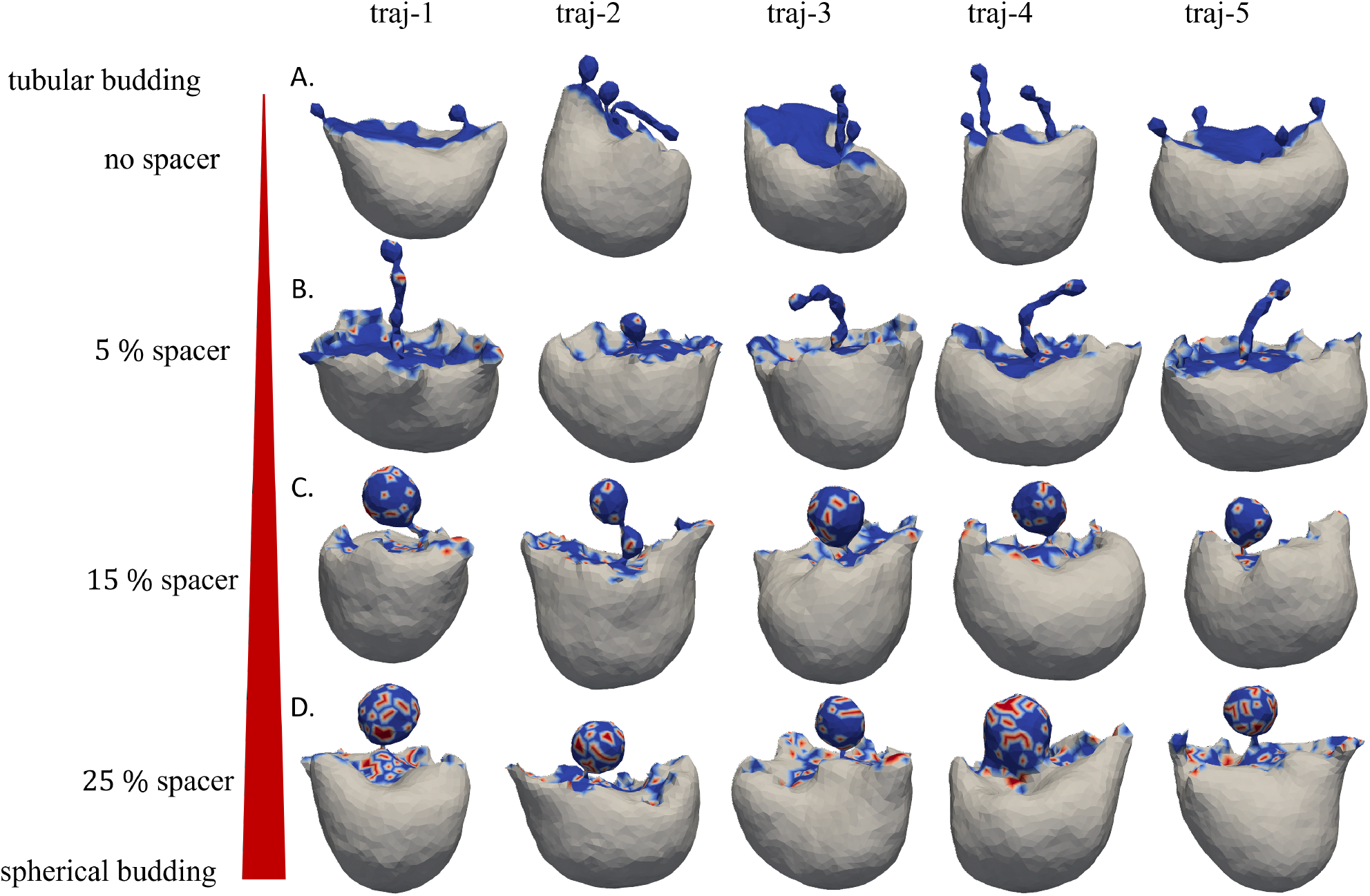
Monte Carlo simulations of triangulated membrane mesh where 20 % vertices are occupied by proteins. The shape of the membrane mesh after 5*×*10^5^ MC steps in 5 parallel runs in the presence of (A) no spacer (B) 5 % (C) 15 % (D) 25 % spacers. Blue regions represent protein and the spacers are depicted with red. The thickness of the red bar is proportional to the number of spacers. The white region is the protein-free membrane surface facing a volume (*K*_*V*_) and area (*K*_*A*_) compressibility of 10 *κ* _*B*_.

In the absence of volume and area compressibility, membrane budding is accompanied by intense deformation of the overall shape because of the propagation of the stress due to the fluid nature of the membrane (Fig-S5). In this case, a larger number of spacers also yields a more spherical membrane bud. To generate membrane budding, uniform distribution of the spacers is crucial. When the spacers are clustered together we do not observe prominent buds (Fig-S6). We also study the effect of anisotropic curvature induction on the shape of membrane budding (Fig-S7). With *c*_||_ = 1 *d*^−1^, *c*_∥_ = 0 *d*^−1^ and *k*_∥_ = *κ*_*B*_, the membrane buds in a tubular fashion. If we turn on *c*_⊥_, multiple branched spheres are generated under a strong coupling limit (*k*_∥*/*⊥_ = *κ*_*B*_). Increasing the value of *c*_∥*/*⊥_ results in a flat shaped budding, which is not consistent with that observed in case of COPII in cells.

## Discussion

This work demonstrates the relationship between the insertion of several amphipathic helices and the creation of specific membrane shapes. This kind of remodeling of the membrane is particularly important in the context of membrane budding triggered by protein coat assemblies.(29) Here, we specifically focus on the case of COPII-mediated membrane budding. Our previous study(15) unraveled a detailed molecular picture of the membrane curvature generation by Sar1, which is known to initiate the COPII-mediated protein trafficking. This study goes beyond the description based on Sar1 and examines the impact of including other inner-coat proteins, Sec23 and Sec24, on the morphology of the membrane budding process. Our findings demonstrate that when Sar1 is densely organized without other COPII proteins, it leads to tubular membrane budding. However, when Sec23 and Sec24 are added, they do not directly contribute to the curvature induction but regulate the surface coverage of Sar1, resulting in a more spherical shape of the bud. Therefore, our results suggest an additional role for Sec23, Sec24, and cargo proteins during COPII transport carrier formation, where they regulate the spacing between Sar1 proteins and thereby facilitate the formation of spherical membrane carriers. This is also supported by recent cryo-EM derived structures showing that Sar1-Sec23-Sec24 trimer units appear as randomly oriented patches on vesicles(24) in contrast to uniformly distributed lattices found on the membrane tubules (5). The amino terminal intrinsically disordered region (IDR) of Sec24, which is not considered in this study, is composed of 70 hydrophilic (13 of which are charged) and 62 hydrophobic residues. It is unclear whether this IDR region helps Sec24 attach to the membrane. However, our simulations indicate that Sec23 and the structured region of Sec24 require Sar1 to maintain their stable attachment with the membrane and thereby form the inner-coat layer. According to our previous studies (15; 19), Sar1 binds to the membrane in both GDP and GTP bound states, but only creates positive membrane curvature in the GTP bound state. Therefore, Sar1 together with Sec23 and Sec24 are able to be associated with the membrane prior to any curvature induction. Afterwards, Sec12 exchanges GDP bound to Sar1 with GTP, which triggers Sar1 to induce curvature on the membrane. However, due to the presence of Sec23, Sec24 and cargo proteins, Sar1 protomers remain scattered on the membrane surface with disordered orientations, leading to spherical budding of the membrane.

Membrane active amphipathic peptides are known to insert and can lead to pore formation.(30; 31) The C-terminal amphipathic helices of complexin have been shown to produce a stable pore in the lipid bilayer when the number of peptides reaches 12.(32) There are just seven hydrophilic residues out of 23 in the amino terminal helix of Sar1. We observe that the amphipathic peptides derived from Sar1 are embedded horizontally to the membrane plane without stretching across despite containing a large amount of hydrophobic residues. The spontaneous transition of a membrane bicelle to a vesicular intermediate triggered by curvature generating protein has been demonstrated previously using the MARTINI model.(33) Here, we establish the dependence of the concentration of amphipathic peptide on the magnitude of curvature induction on membrane bicelle (Fig-2). We also reveal that the relative membrane sculpting efficiency of amphipathic peptides decreases in the absence of the entire protein segment. In the case of BAR(34; 35) domains and the ESCRT machinery(36), proteins form an intrinsically curved filament, which is key to the process of membrane bending by these proteins. On the contrary, the lack of dependency of the membrane curvature induction on the inter-peptide angle indicates that the curvature induction is isotropic in the case of COPII. Our DTS simulations also reveal that isotropic curvature induction on the protein-bound vertices is essential to producing spherical membrane bud at optimal spacer concentration.

The conventional mechanism of cargo transport through COPII-coated membrane vesicles has been challenged by two recent experiments.(37; 38; 39) These results suggest that COPII localizes at the neck of the membrane bud, defining a boundary between ER and ER-Golgi intermediate compartments (ERGIC). Further, the inhibition of Sar1 reduced the recruitment of Sec23 on the ER membrane and thus disrupted the formation of a proper COPII assembly.(38) Our MARTINI based simulations also support this finding by showing that Sar1 recruits Sec23-Sec24 to form the inner-coat layer. However, the mechanism by which such a ring-shaped COPII collar can produce a membrane bud remains unclear.

In summary, we offer a mechanistic overview of the complex interplay between multiple proteins from COPII family in regulating the shape of the coated membrane surface. We cover a broad range of length scales by employing atomistic, MARTINI, and Helfrich Hamiltonian based mesoscale simulations to establish the role of spacer proteins in producing spherical membrane buds. Here, we focus our study on the inner-coat proteins Sar1, Sec23, and Sec24, which are considered to be the key players in the membrane budding process at the subdomains of ER. In addition to the inner-coat layer, Sec13 and Sec31 form the outer-coat layer, which has a cage-like structure and promotes vesicle fission.(40) It is not clear whether the highly bent outer-coat layer also contributes to the curvature induction and membrane budding process. An intriguing possibility is that interactions between the inner and outer coat protein facilitate appropriate spacing between Sar1 protomers to drive the budding of spherical transport carriers. Although our approach of modeling the inner-coat layer is sufficient to explain the shapes of remodeled membranes in both *in vitro* and *in vivo* conditions, it is important in the next step to directly consider the roles of Sec13 and Sec31 in this process.

## Methods

### Atomistic simulations

We follow the protocol described in Paul et. al. (15) to perform the atomistic membrane simulations. To estimate the membrane penetration depth of the peptide dimer, we perform simulations in the presence of a continuous membrane that interacts with its periodic images along *X* and *Y* dimensions. We start with the h-GTP dimer model of Sar1 inserted in an atomistic lipid bilayer (10*nm×*10*nm*) where the composition of the bilayer is 66 % DOPC, 21 % DOPE, 8 % DOPS, and 5 % DOPA; i.e., the membrane is 13 % anionic. We consider only the amino-terminal amphipathic helix (residue 1-23) embedded in the membrane bilayer, discarding the rest of the protein and solvent. We then re-solvate and re-ionize the system to make it charge-neutral and to maintain the physiological (0.15 M) salt concentration. The solvated system is then energy minimized followed by NVT equilibration in which a harmonic restraint is initially applied to all the heavy atoms of the peptides and membrane and then gradually released during equilibration. Subsequently, NPT simulations are carried out for 160 ns with the Nosé-Hoover thermostat and Parinello-Rahman barostat (semiisotropic pressure coupling) to control the temperature and pressure of the system. All simulations are performed using GROMACS(41; 42) version 2018.3 and the CHARMM36m(43) force field with the TIP3P explicit solvent model. The h-GTP dimer showed ∼40 % penetration depth (Fig-S11 A in the SI of Ref. (15)) inside the membrane where 100 % represents half of the membrane thickness (distance between the phosphate plane and the center of the membrane) as defined by Paul et. al.(15). In the absence of the rest of the protein, the peptides exhibit a deeper insertion depth (∼65 %). We quantify the magnitude of the partitioning of the hydrophobic/hydrophilic residues of the amphipathic helices at the membrane-water interface by a quantity *µ*, which is defined as follows,

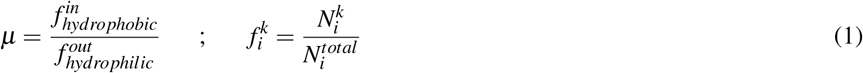

where *I* ∈ [*hydrophobic*; *hydrophilic*] and *k*∈ [*in*; *out*]. *i* is the type of residue the atom belongs and *k* is the state of the atom with respect to the membrane plane (outside/inside the membrane). 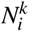 is the number of atoms of type *i* and state *k* and 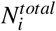 is the total number of atoms of type *i* present in the helix.

We model the bicelle system by replicating the continuous membrane system along +*X* : −*X* and +*Y* : −*Y* directions using the gmx genbox utility followed by the deletion of membrane segments near the edges to break the membrane continuity along *X* and *Y*. As a result, the membrane size is 19*nm×*19*nm* whereas the size of the simulation box is 22*nm×*22*nm* along the *X* and *Y* dimensions. We employ anisotropic pressure coupling while keeping the other parameters the same as above. To avoid rotation of the bicelle, four lipid atoms at the periphery of the bicelle are restrained for their movement along the *Z* axis. PLUMED(44) version 2.5.3(45) is used to restrain the inter-peptide angle to *π* in arrangement-2. VMD version 1.9.3(46) is used for visualization and analysis. The helical wheel diagram is constructed using NetWheels(47).

### Coarse Grained MARTINI simulation

We perform explicit solvent coarse grained MARTINI simulations with Sar1 (Fig-S3) and multiple units of the Sar1-Sec23-Sec24 trimer on a lipid bilayer. We convert the atomistic models of Sar1-Sec23-Sec24 (PDB code: 6GNI(5)) trimer into MARTINI3.0(48) model using martinize2(49) with the elastic bond force constant of 1300 kJ/mol/*nm*^2^ and a decay factor 0.8. To yield efficient insertion of the amphipathic helix of Sar1 inside the membrane bilayer, we remove all the elastic bonds from its amino terminus (residue 1-23). We do not consider the amino-terminal loop region (residue 1-132) of Sec24 in our study to gain computational efficiency. After building the MARTINI model of the proteins, we construct a 40 ∼ *nm×*30 *nm* membrane bilayer with 87 % DOPC and 13 % DOPS lipid molecules and place the protein molecules on top using the insane tool. Then, we add water beads and *Na*^+^*/Cl*^−^ to neutralize the system and maintain a physiological salt concentration. The simulation box size is ∼ 40 *nm* × 30 *nm×*30 *nm*. Then the system is energy minimized followed by short (∼1 *ns*) NPT equilibration with Berendsen thermostat and barostat (*τ*_*P*_ 12 *ps*^−1^). Finally, a ∼2− 5 *µs* production run is carried out with the equilibrated configuration. In this step, V-rescale thermostat and Parrienllo-Rahman barostat are used with the same *τ*_*P*_. The time step in the MARTINI simulations is 20 fs.

### Dynamic Triangulated Surface simulation

We carry out the Monte Carlo (MC) simulation of the Dynamic Triangulated Surface (DTS) model following the strategy described by Ramakrishnan et. al.(21; 50) In this simulation the membrane dynamics is governed by the Helfrich Hamiltonian at the mesoscopic length scale where the membrane is considered as a surface discretized by triangles. The presence of curvature-inducing proteins is represented by vertices that have intrinsic non-zero curvatures. Here, instead of using the formalism developed by Ramakrishnan et al., we utilize isotropic curvature induction as discussed by Pezeshkian et. al.(27).

Therefore, the Hamiltonian in our case is

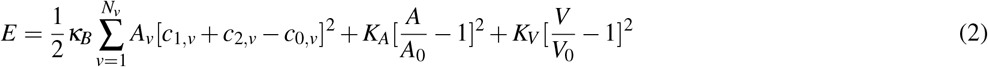

where *κ*_*B*_ is the membrane bending modulus, which is taken to be 20 *k*_*B*_*T, v* is the vertex index, *A*_*v*_ is the total area of all the neighboring triangles of vertex *v, c*_1,*v*_ and *c*_2,*v*_ are the principal curvatures, and *c*_0,*v*_ is the intrinsic curvature at vertex *v. c*_0,*v*_ is 0 *d*^−1^ for the membrane and spacer containing vertex while 1 *d*^−1^ for the protein containing vertex. *K*_*A*_ and *K*_*V*_ are the area and volume compressibilities, respectively and we choose their value as 10 *κ*_*B*_ for the non-protein vertices to avoid significant deformation of the membrane. *A* is the total area of the membrane during the MC run and *A*_0_ is the initial value. *V* and *V*_0_ are defined in a similar way for the total volume. *d* is the unit length in the mesoscopic scale. By comparing the curvature generated in the atomistic and mesoscopic length scales, we estimate the value of *d* ∼10 *nm*. Therefore, one node (vertex) in the DTS membrane is equivalent to the size of a membrane bicelle that is simulated at the atomistic scale. Vertex movement and link flips are considered as the MC moves. We allow nematic exchange between protein (blue) and spacers (red) while no nematic exchange is allowed between membrane and protein/spacer. A total of 5*×*10^5^ MC steps are carried out. For comparison, we also perform DTS simulations using anisotropic curvature induction condition as follows,

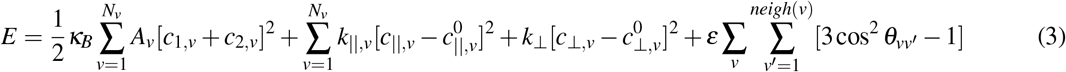

Where

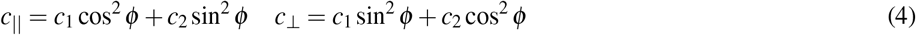

Here *ϕ* is the angle between the nematic vector and the principal curvature at the vertex. *θ* is the angle between the nematic vectors in the tangent plane at the neighboring vertices. With *k*_∥_ and *k*_⊥_, curvature induction can be separately coupled along the parallel and perpendicular directions to the nematic vector. The total number of vertices in the membrane vesicle is 2030, 20% of which are occupied by proteins (Sar1). Among the protein-containing vertices, we place spacers representing Sec23/Sec24 with varying amounts (5-75 %). To induce aggregation among the spacers, we use *ε* within the isotropic curvature induction framework where the Hamiltonian looks like the following.

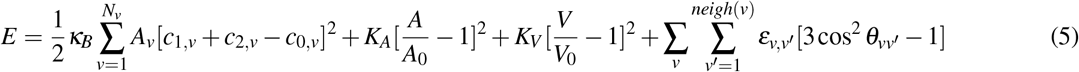

where *ε*_spacer,spacer_ = 3 *K*_B_*T, ε*_protein,spacer_ = 0 *K*_B_*T* and *ε*_protein,protein_ = 0 *K*_B_*T*

We use paraview(51) to visualize the membrane in the form of the triangulated mesh where surface representation is selected and the colour scheme is based on the phases of the vertices. Non-protein containing vertices (grey) belong to phase 2 whereas proteins (blue) and spacers (red) containing vertices belong to phase 1 and 3 respectively.

## Acknowledgements

The work is supported in part by the NSF grant to QC (CHE-2154804). This work used Delta at the National Center for Supercomputing Application (NCSA) through allocation MCB110014 from the Advanced Cyberinfrastructure Coordination Ecosystem: Services & Support (ACCESS) program(52), which is supported by National Science Foundation grants #2138259, #2138286, #2138307, #2137603, and #2138296. A part of the computational work was performed on the Shared Computing Cluster which is administered by Boston University’s Research Computing Services (URL: www.bu.edu/tech/support/research/). We thank Dr. Ramakrishnan Natesan for the discussion regarding the mesoscale model of membranes. We also thank Dr. Xiao-Han Li for useful comments on the manuscript.

## Author contributions statement

S.P., A.A., and Q.C. designed research; S.P. performed research; S.P. and Q.C. analyzed data; and S.P., A.A., and Q.C. wrote the paper.

## References

1. Hughes, H. & Stephens, D. J. Assembly, organization, and function of the copii coat. Histochem. cell biology 129, 129–151 (2008).

2. d’Enfert, C., Wuestehube, L. J., Lila, T. & Schekman, R. Sec12p-dependent membrane binding of the small gtp-binding protein sar1p promotes formation of transport vesicles from the er. J. Cell Biol. 114, 663–670 (1991).

3. Lee, M. C. et al. Sar1p n-terminal helix initiates membrane curvature and completes the fission of a copii vesicle. Cell 122, 605–617 (2005).

4. Bi, X., Corpina, R. A. & Goldberg, J. Structure of the sec23/24–sar1 pre-budding complex of the copii vesicle coat. Nature 419, 271–277 (2002).

5. Hutchings, J., Stancheva, V., Miller, E. A. & Zanetti, G. Subtomogram averaging of copii assemblies reveals how coat organization dictates membrane shape. Nat. Commun. 9, 1–8 (2018).

6. Shaywitz, D. A., Espenshade, P. J., Gimeno, R. E. & Kaiser, C. A. Copii subunit interactions in the assembly of the vesicle coat. J. Biol. Chem. 272, 25413–25416 (1997).

7. Bi, X., Mancias, J. D. & Goldberg, J. Insights into copii coat nucleation from the structure of sec23^*^ sar1 complexed with the active fragment of sec31. Dev. cell 13, 635–645 (2007).

8. Hutchings, J. et al. Structure of the complete, membrane-assembled copii coat reveals a complex interaction network. Nat. communications 12, 2034 (2021).

9. Matsuoka, K. et al. Copii-coated vesicle formation reconstituted with purified coat proteins and chemically defined liposomes. Cell 93, 263–275 (1998).

10. Raote, I. et al. Tango1 builds a machine for collagen export by recruiting and spatially organizing copii, tethers and membranes. Elife 7, e32723 (2018).

11. Raote, I. et al. A physical mechanism of tango1-mediated bulky cargo export. Elife 9, e59426 (2020).

12. Long, K. R. et al. Sar1 assembly regulates membrane constriction and er export. J. Cell Biol. 190, 115–128 (2010).

13. Hariri, H., Bhattacharya, N., Johnson, K., Noble, A. J. & Stagg, S. M. Insights into the mechanisms of membrane curvature and vesicle scission by the small gtpase sar1 in the early secretory pathway. J. molecular biology 426, 3811–3826 (2014).

14. Kasberg, W., Luong, P., Swift, K. A. & Audhya, A., Nutrient deprivation alters the rate of copii subunit recruitment at er subdomains to tune secretory protein transport. Nat. Commun. 14, 8140 (2023).

15. Paul, S., Audhya, A. & Cui, Q. Molecular mechanism of gtp binding-and dimerization-induced enhancement of sar1-mediated membrane remodeling. Proc. Natl. Acad. Sci. 120, e2212513120 (2023).

16. Yoshihisa, T., Barlowe, C. & Schekman, R. Requirement for a gtpase-activating protein in vesicle budding from the endoplasmic reticulum. Science 259, 1466–1468 (1993).

17. Wendeler, M. W., Paccaud, J.-P., & Hauri, H.-P. Role of sec24 isoforms in selective export of membrane proteins from the endoplasmic reticulum. EMBO reports 8, 258–264 (2007).

18. Kasberg, W. et al. The sar1 gtpase is dispensable for copii-dependent cargo export from the er. Cell Reports 42 (2023).

19. Hanna, M. G. et al. Sar1 gtpase activity is regulated by membrane curvature. J. Biol. Chem. 291, 1014–1027 (2016).

20. Ramakrishnan, N., Ipsen, J. H. & Kumar, P. S. Role of disclinations in determining the morphology of deformable fluid interfaces. Soft Matter 8, 3058–3061 (2012).

21. Ramakrishnan, N., Kumar, P. S. & Radhakrishnan, R. Mesoscale computational studies of membrane bilayer remodeling by curvature-inducing proteins. Phys. reports 543, 1–60 (2014).

22. Mandal, T., Spagnolie, S. E., Audhya, A. & Cui, Q. Protein-induced membrane curvature in coarse-grained simulations. Biophys. J. 120, 3211–3221 (2021).

23. Mondal, S. & Cui, Q. Coacervation of poly-electrolytes in the presence of lipid bilayers: mutual alteration of structure and morphology. Chem. Sci. 13, 7933–7946 (2022).

24. Pyle, E. & Zanetti, G. Cryo-electron tomography reveals how copii assembles on cargo-containing membranes. bioRxiv 2024–01 (2024).

25. Ramakrishnan, N., Kumar, P. S. & Ipsen, J. H. Membrane-mediated aggregation of curvature-inducing nematogens and membrane tubulation. Biophys. journal 104, 1018–1028 (2013).

26. Kumar, G., Ramakrishnan, N. & Sain, A. Tubulation pattern of membrane vesicles coated with biofilaments. Phys. Rev. E 99, 022414 (2019).

27. Pezeshkian, W. & Ipsen, J. H. Fluctuations and conformational stability of a membrane patch with curvature inducing inclusions. Soft matter 15, 9974–9981 (2019).

28. Pezeshkian, W. & Ipsen, J. H. Mesoscale simulation of biomembranes with freedts. Nat. Commun. 15, 548 (2024).

29. Stachowiak, J. C., Brodsky, F. M. & Miller, E. A. A cost–benefit analysis of the physical mechanisms of membrane curvature. Nat. cell biology 15, 1019–1027 (2013).

30. Kabelka, I. & Vácha, R. Advances in molecular understanding of α-helical membrane-active peptides. Accounts Chem. Res. 54, 2196–2204 (2021).

31. Ulmschneider, J. P. & Ulmschneider, M. B. Molecular dynamics simulations are redefining our view of peptides interacting with biological membranes. Accounts chemical research 51, 1106–1116 (2018).

32. Courtney, K. C. et al. The complexin c-terminal amphipathic helix stabilizes the fusion pore open state by sculpting membranes. Nat. structural & molecular biology 29, 97–107 (2022).

33. Bhaskara, R. M. et al. Curvature induction and membrane remodeling by fam134b reticulon homology domain assist selective er-phagy. Nat. communications 10, 2370 (2019).

34. Arkhipov, A., Yin, Y. & Schulten, K. Four-scale description of membrane sculpting by bar domains. Biophys. J. 95, 2806–2821 (2008).

35. Arkhipov, A., Yin, Y. & Schulten, K. Membrane-bending mechanism of amphiphysin n-bar domains. Biophys. J. 97, 2727–2735 (2009).

36. Mandal, T., Lough, W., Spagnolie, S. E., Audhya, A. & Cui, Q. Molecular simulation of mechanical properties and membrane activities of the escrt-iii complexes. Biophys. J. 118, 1333–1343 (2020).

37. Shomron, O. et al. Copii collar defines the boundary between er and er exit site and does not coat cargo containers. J. Cell Biol. 220, e201907224 (2021).

38. Weigel, A. V. et al. Er-to-golgi protein delivery through an interwoven, tubular network extending from er. Cell 184, 2412–2429 (2021).

39. Malis, Y., Hirschberg, K. & Kaether, C. Hanging the coat on a collar: Same function but different localization and mechanism for copii. BioEssays 44, 2200064 (2022).

40. Zanetti, G., Pahuja, K. B., Studer, S., Shim, S. & Schekman, R. Copii and the regulation of protein sorting in mammals. Nat. cell biology 14, 20–28 (2012).

41. Abraham, M. J. et al. Gromacs: High performance molecular simulations through multi-level parallelism from laptops to supercomputers. SoftwareX 1, 19–25 (2015).

42. Berendsen, H. J., van--der--Spoel, D. & van--Drunen, R. Gromacs: A message-passing parallel molecular dynamics implementation. Comput. Phys. Commun. 91, 43–56 (1995).

43. Huang, J. et al. Charmm36m: an improved force field for folded and intrinsically disordered proteins. Nat. Methods 14, 71–73 (2017).

44. Promoting transparency and reproducibility in enhanced molecular simulations. Nat. methods 16, 670–673 (2019).

45. Tribello, G. A., Bonomi, M., Branduardi, D., Camilloni, C. & Bussi, G. Plumed 2: New feathers for an old bird. Comput. physics communications 185, 604–613 (2014).

46. Humphrey, W., Dalke, A. & Schulten, K. Vmd: visual molecular dynamics. J. molecular graphics 14, 33–38 (1996).

47. Mol, A. R., Castro, M. S. & Fontes, W. Netwheels: A web application to create high quality peptide helical wheel and net projections. BioRxiv 416347 (2018).

48. Souza, P. C. et al. Martini 3: a general purpose force field for coarse-grained molecular dynamics. Nat. methods 18, 382–388 (2021).

49. Kroon, P. C. et al. Martinize2 and vermouth: Unified framework for topology generation. arXiv preprint arXiv:2212.01191(2022).

50. Ramakrishnan, N., Kumar, P. S. & Ipsen, J. H. Monte carlo simulations of fluid vesicles with in-plane orientational ordering. Phys. Rev. E 81, 041922 (2010).

51. Ahrens, J., Geveci, B., Law, C., Hansen, C. & Johnson, C. 36-paraview: An end-user tool for large-data visualization. The visualization handbook 717, 50038–1 (2005).

52. Boerner, T. J., Deems, S., Furlani, T. R., Knuth, S. L. & Towns, J. ACCESS: Advancing innovation: NSF’s advanced cyberinfrastructure coordination ecosystem: Services & support. In Practice and Experience in Advanced Research Computing (PEARC ‘23) (2023).

