## Supporting Information for "Delineating the shape of COPII coated membrane bud"

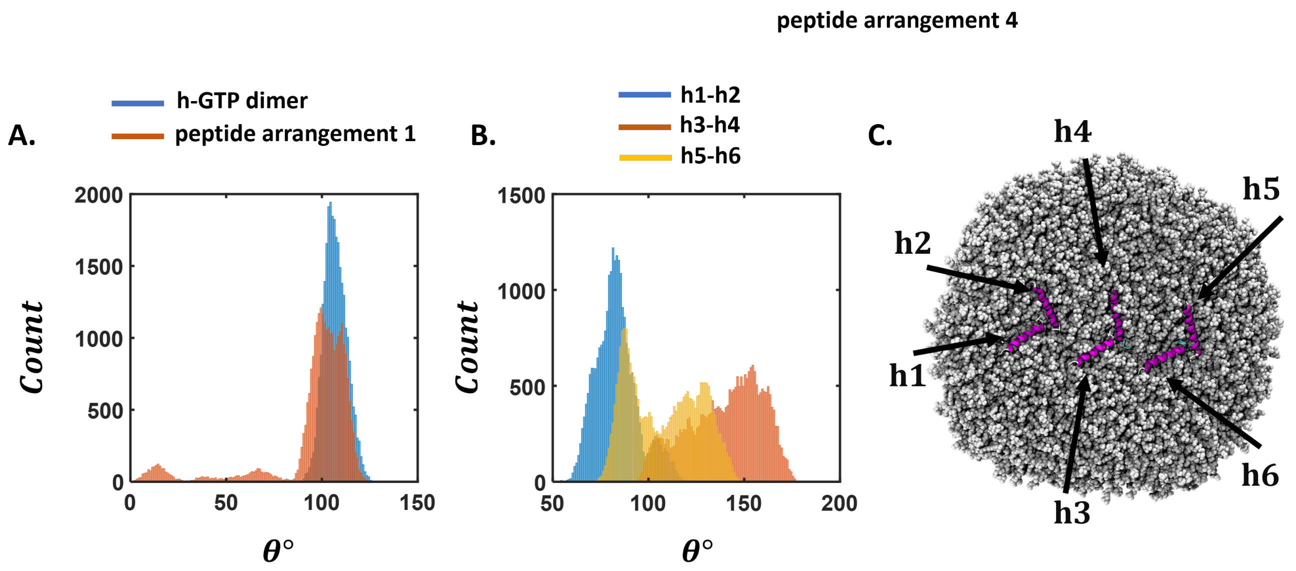


**Figure S1.** The relative orientation of the helical amphipathic peptides in membrane bicelle simulation. (A) Distribution of the inter helix angle in h-GTP dimer and peptide arrangement 1 during membrane bicelle simulation. In the presence of the entire protein segment, the angle between the amphipathic amino-terminal helix exhibits a sharp distribution whereas when the peptides are present alone the distribution becomes skewed. (B) A similar plot in the case of peptide arrangement 4 where 3 pairs of helices are present. The angle between h1-h2 shows a sharper distribution compared to the cases of the other two helix pairs. (C) Depiction of the helices on the membrane bicelle in the case of peptide arrangement 4. Overall, we observe a broad range of distributions of the inter-helix angles when the peptides are present in the absence of the rest of the protein segment.

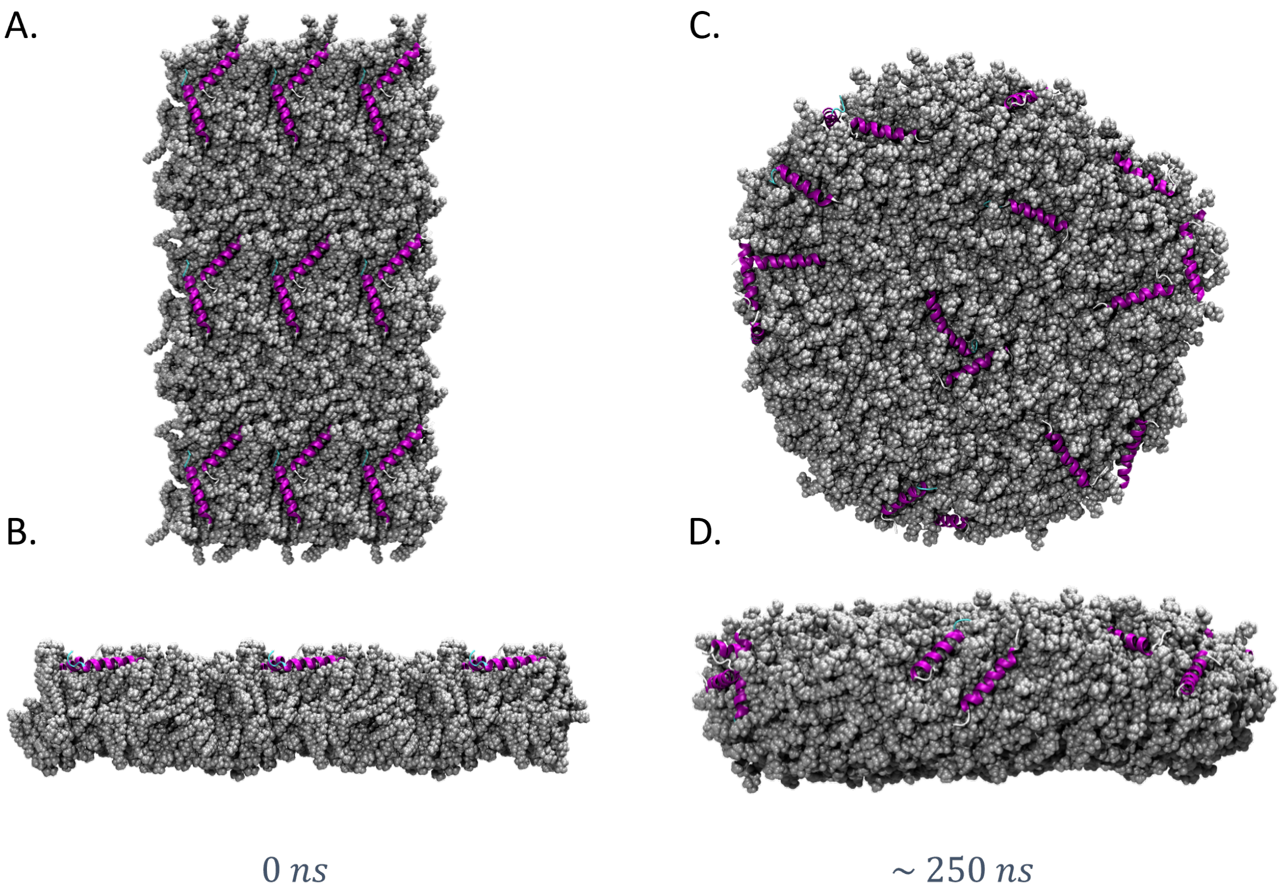


**Figure S2.** Complete coverage of one face of the membrane bicelle by amphipathic peptides does not lead to any membrane curvature induction. Top and Side view of snapshot of membrane bicelle at (A-B) 0 *ns* and (C-D) 250 *ns* where a total of 18 amphipathic peptides derived from the amino terminal amphipathic helix of Sar1 is placed on one side. Although the peptide number is higher in this case compared to that in the case of arrangement-4, we do not observe any membrane bending. Instead, some of the peptides translocate to the positively curved edge region, reflecting the curvature-sensing property of the peptides. This observation indicates that when amphipathic peptides occupy the edge region of the membrane bicelle, inter-leaflet stress is not sufficient to induce membrane curvature.

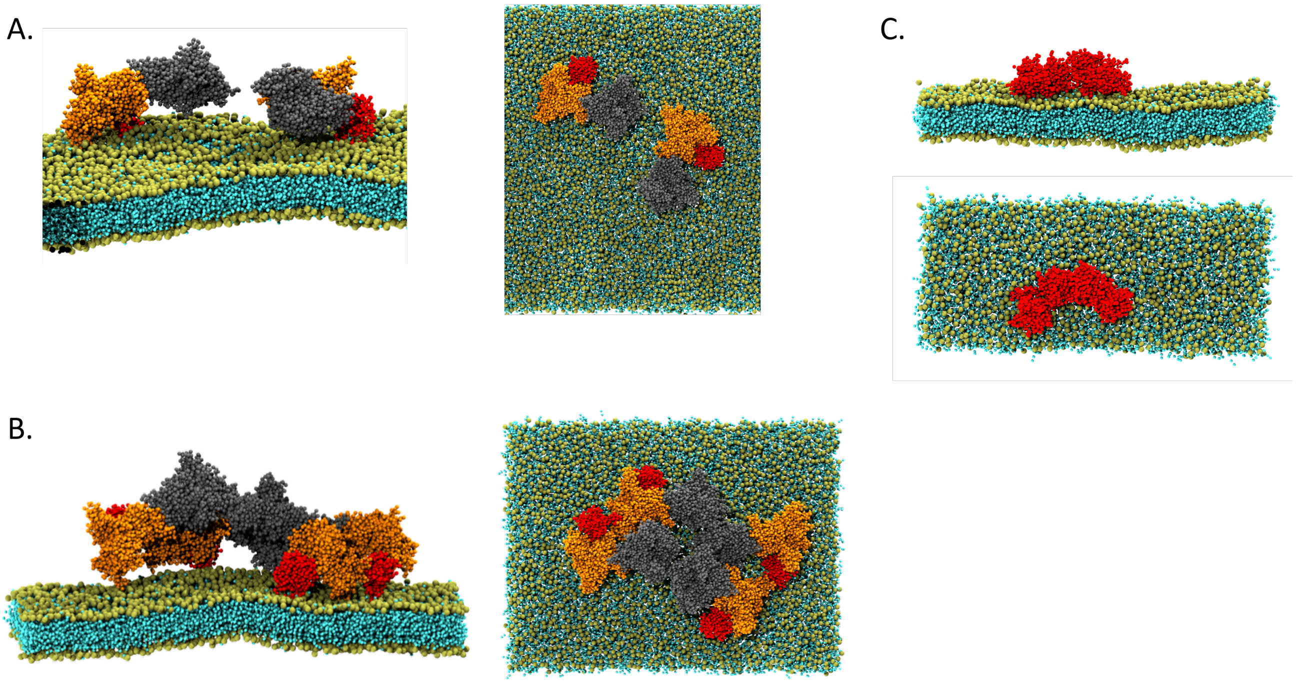


**Figure S3.** Coassembly of Sar1-Sec23-Sec24 on the membrane using the MARTINI model. Side and top view of (A) two and (B) four Sar1-Sec23-Sec24 trimers on the membrane. Sar1 is colored in red, Sec23 in orange, and Sec24 in gray. In the latter case, one out of four Sar1 proteins remains out of the membrane plane due to crowding. (C) Four Sar1 proteins remain attached in the absence of Sec23 and Sec24, producing a near linear assembly. $d_{amino}$ is computed in the case of B vs. C in Fig- 3 in the main text.

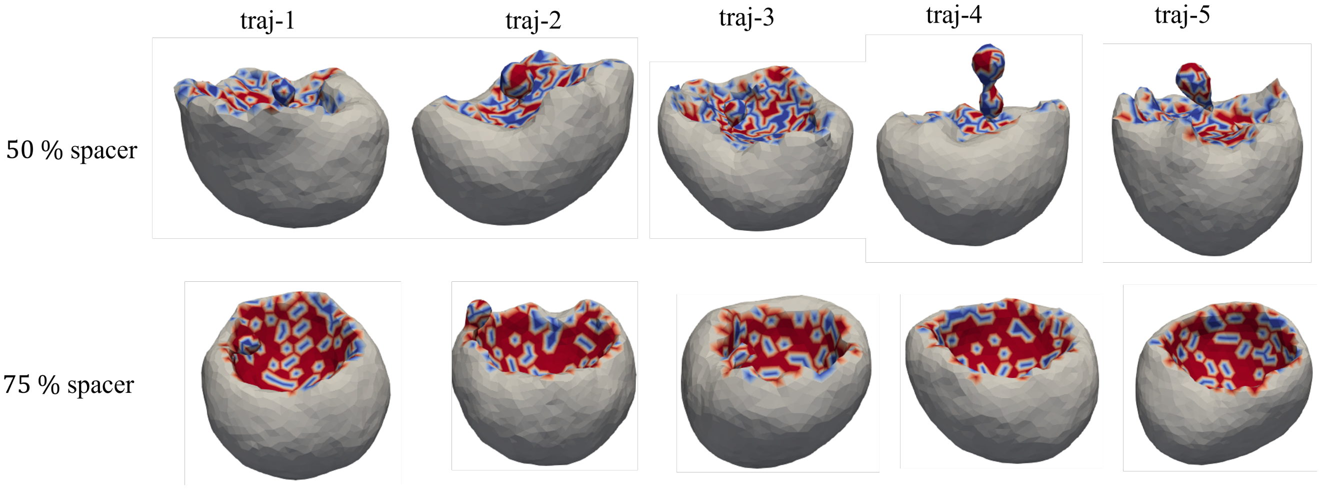


**Figure S4.** Membrane surface as a triangulated mesh after $5 \times 10^5$ MC steps in five independent MC runs when 50 % and 75 % of the protein-containing vertices are occupied by spacers. Blue regions are protein-containing vertices and red represents the spacers. The rest of the vertices represent the protein-free membrane surface. With the 50 % spacer coverage much smaller spherical budding is observed compared to that in the case of 25 % (Fig-4). When the spacer quantity reaches 75 % no prominent budding is observed. Instead, the condition yields a small amount of negative curvature due to the lack of volume and area compressibility of the spacer-containing region of the membrane.

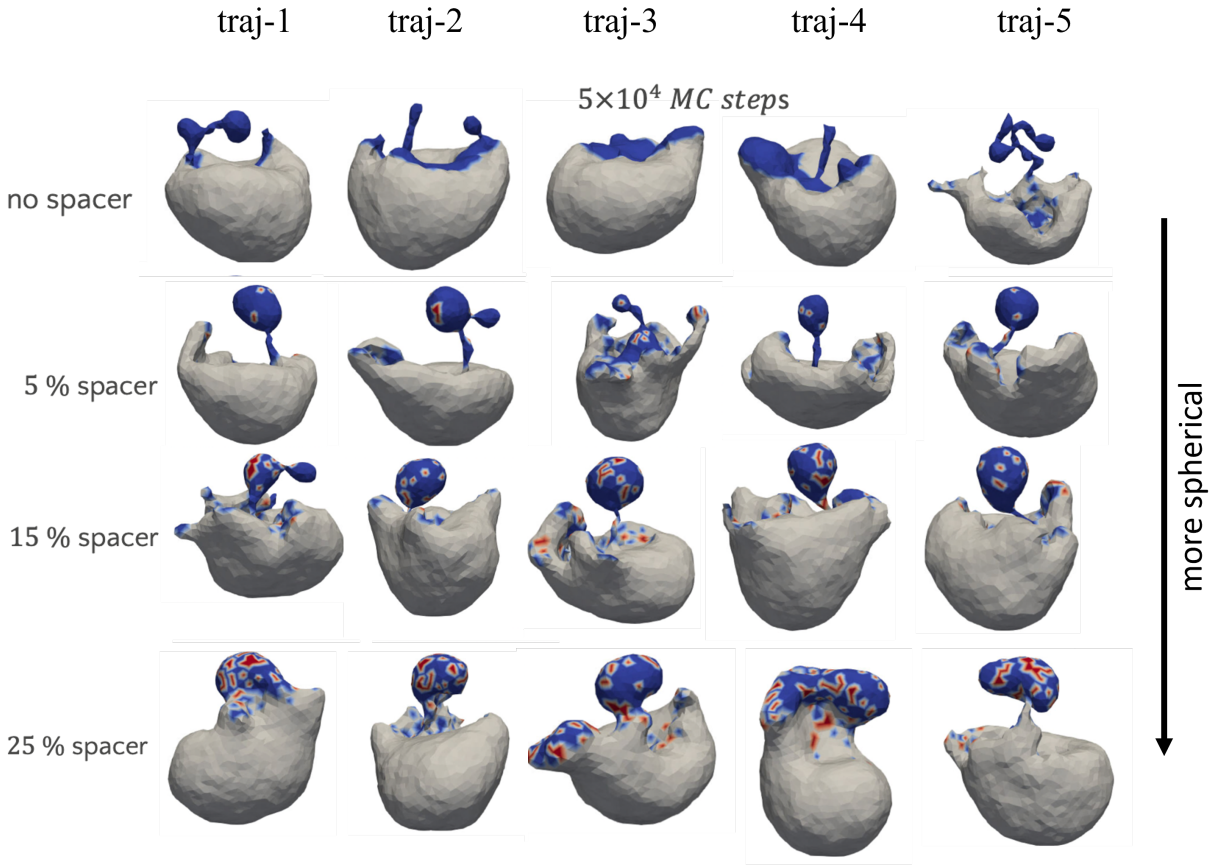


**Figure S5.** Effect of spacers on the shape of the membrane buds when no area and volume compressibility are applied on any vertices. Here, we again demonstrate that with the increasing amount of spacers, the tubular shape of the membrane buds transforms into a more spherical shape (as also shown in Fig-4). However, in contrast to Fig-4, here we observe more intense deformation of the overall shape of the membrane, especially near the boundary of the protein-containing region as $K_A = K_V = 0$. All the snapshots are captured after $5 \times 10^4$ MC steps. @@Do we really need traj-6 while other cases only contain 5 snapshots?@@

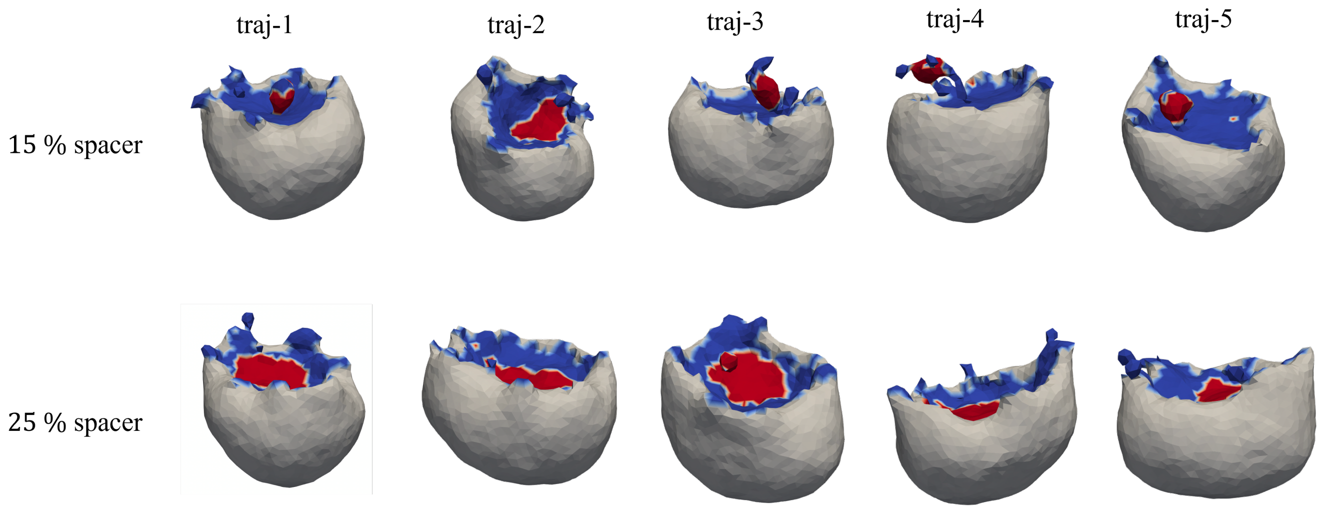


**Figure S6.** Aggregated spacers are unable to generate spherical membrane budding. Snapshots of membrane surface where 20 % vertices are occupied by proteins and 15 – 25 % of the protein containing vertices are occupied by non-curvature inducing spacers. Spacers are clustered due to the $\varepsilon$ term in Eqn. 5. $K_A$ and $K_V$ are set to 10 $\kappa_B$ for the membrane vertices similar to that in the case of Fig-4. Tiny membrane buds are visible in the case of 15 % spacer whereas no membrane budding is observed in the case of 20 % spacer. On the other hand, when spacers are distributed uniformly among the protein-containing vertices, spherical membrane budding is observed in both cases as shown in Fig-4.

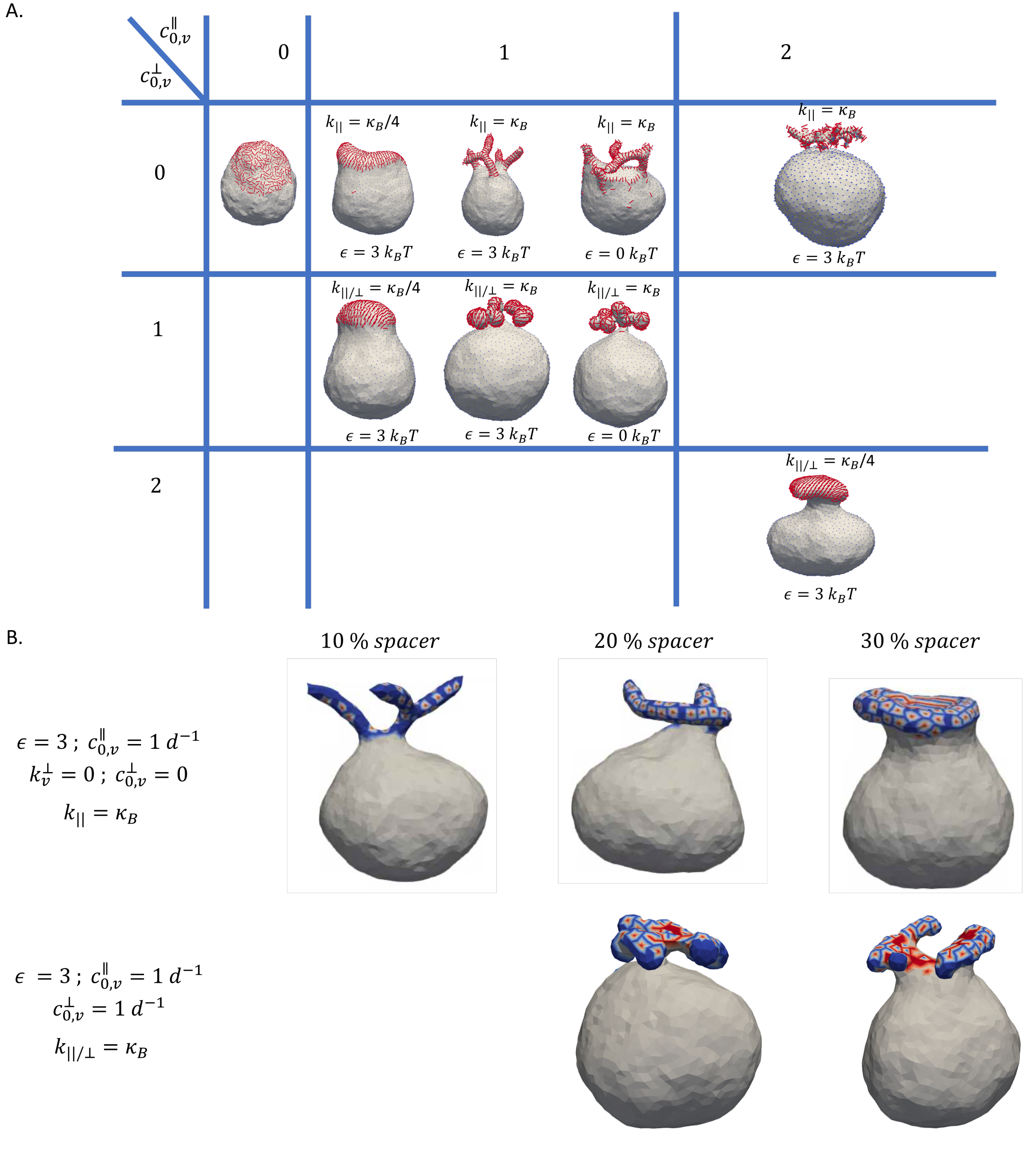


**Figure S7.** (A) The shape of the membrane bud is presented in a tabular format when proteins (nematics shown in red line) are anisotropically coupled as described in Eqn. 3-4. We provide snapshots of the membrane with 20 % vertices occupied by proteins under different values of $c_{0,v}^{\parallel}$, $c_{0,v}^{\perp}$, $k_{\parallel}$, and $k_{\perp}$. $5 \times 10^5$ MC steps are conducted using the Hamiltonian shown in Eqn. 3. When $c_{0,v}^{\perp} = 0$ but $c_{0,v}^{\parallel} \neq 0$, tubular budding is observed when $k_{\parallel} = \kappa_B$. Under the condition of $c_{0,v}^{\perp} = c_{0,v}^{\parallel} = 1\, d^{-1}$, branched spherical budding is generated when $k_{\parallel} = \kappa_B$. The shape of the membrane bud becomes flat when $c_{0,v}^{\perp} = c_{0,v}^{\parallel} = 2\, d^{-1}$ and $k_{\parallel} = \kappa_B/4$. The value of $\varepsilon$ does not impact the shape of the membrane bud much. (B) Effect of spacer on the topology of the membrane budding under the anisotropic curvature induction condition. Color coding is the same as Fig-4 and S4-S6. As the spacer content increases, the membrane bud adopts a flat disc-like shape which is not commonly observed in experiments. Therefore, spacer induced sphericity of the membrane bud only takes place when curvature induction occurs in an isotropic manner as described in Eqn. 2.
